# Estimation of sinking velocities using free-falling dynamically scaled models: foraminifera as a test case

**DOI:** 10.1101/2020.06.18.159780

**Authors:** Matthew Walker, Stuart Humphries, Rudi Schuech

## Abstract

The velocity of settling particles is an important determinant of distribution in extinct and extant species with passive dispersal mechanisms, such as plants, corals, and phytoplankton. Here we adapt dynamic scaling, borrowed from engineering, to determine settling velocities. Dynamic scaling leverages physical models with relevant dimensionless numbers matched to achieve similar dynamics to the original object. Previous studies have used flumes, wind tunnels, or towed models to examine fluid flows around objects with known velocities. Our novel application uses free-falling models to determine the unknown sinking velocities of planktonic foraminifera – organisms important to our understanding of the Earth’s current and historic climate. Using enlarged 3D printed models of microscopic foraminifera tests, sunk in viscous mineral oil to match their Reynolds numbers and drag coefficients, we predict sinking velocities of real tests in seawater. This method can be applied to study other settling particles such as plankton, spores, or seeds.

**Summary Statement:** We developed a novel method to determine the sinking velocities of biologically important microscale particles using 3D printed scale models.

## Introduction

The transport of organisms and biologically derived particles through fluid environments strongly influences their spatiotemporal distributions and ecology. In up to a third of terrestrial plants (Willson et al., 1990), reproduction is achieved through passive movement of propagules (e.g., seeds) on the wind. In aquatic environments, propagules of many sessile groups from corals (Jones et al., 2015) to bivalves (Booth, 1983) are dispersed by ambient currents, eventually settling out of the water column to their final locations. Furthermore, most dead aquatic organisms (from diatoms to whales) sink, transporting nutrients to deeper water and contributing to long term storage of carbon (De La Rocha & Passow, 2007). In the case of microfossils, sinking dynamics of the original organisms even influences our reconstructions of the Earth’s paleoclimate (Van Sebille et al., 2015). Crucially, the horizontal distances over which all these biological entities are transported, and therefore their distributions, are affected by their settling velocities (Ali et al., 2011).

Measuring the individual settling velocities of small particles directly is challenging, especially when they are too small to be imaged easily without magnification (e.g. Walsby and Holland, 2006). Here we apply dynamic scaling, an approach commonly used in engineering, to circumvent this difficulty and accurately quantify the kinematics of sub-millimetre scale free-falling particles using enlarged physical models and standard high-definition cameras. We use scaled-up physical models in a high-viscosity fluid, enabling easy measurements of settling speed, orientation, and other parameters using inexpensive standard high-definition web cameras. While dynamically scaled models have previously been employed to study a number of problems in biological fluid mechanics (e.g. Vogel, Ellington and Kilgore, 1973; Vogel, 1987; Vogel, 1994; Koehl, 2003), the study of freely-falling particles of complex shape – for which settling speed is the key unknown parameter – presents a unique challenge to experimental design that we overcome in this work.

Engineering problems such as aircraft and submarine design often are approached using scaled-down models in wind tunnels or flumes to examine fluid flows around the model and the resulting fluid dynamic forces it is subjected to. To ensure that the behaviour of the model system is an accurate representation of real life, similarity of relevant physical phenomena must be maintained between the two. If certain dimensionless numbers (i.e., ratios of physical quantities such that all dimensional units cancel) that describe the system are equal between the life-size original and the scaled-down model, “similitude” is achieved and all parameters of interest (e.g., velocities and forces) will be proportional between prototype and model (Zohuri, 2015). Intuitively, the model and real object must be geometrically similar (i.e., have the same shape), so that the dimensionless ratio of any length between model and original, *Length*_*model*_/*Length*_*real*_, is constant – this is the scale factor (*S*) of the model. Less obvious is the additional requirement of dynamic similarity, signifying that the ratios of all relevant forces are constant. For completely immersed objects moving steadily through the surrounding fluid, dynamic similarity is achieved by matching the Reynolds number (*Re*).

*Re* is a measure of the ratio of inertial to viscous forces in the flow (Batchelor, 2000; within a biological context Vogel, 1994), and is typically defined as:

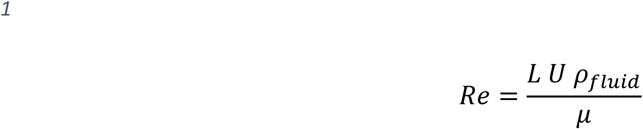

where *ρ*_*fluid*_ is density of the fluid (kg m^-3^); *L* is a characteristic length (m) of the object; *U* is the object’s velocity (m s^-1^); and *μ* is the dynamic viscosity (*N s m*^-2^, or *pa s*) of the fluid. In cases where *L U ρ*_*fluid*_ is large compared to *μ*, e.g. fish, birds, and whales (*Re* ≈ 3×10^6^ - 3×10^9^, Vogel, 1994), inertial forces dominate. In cases where *L U ρ*_*fluid*_ is relatively small compared to *µ*, e.g. bacteria, sperm (Re ≈ 1×10^−5^ - 3×10^−2^), viscous forces dominate. Finally, when *L U ρ*_*fluid*_ is of comparable magnitude to *μ, Re* is intermediate and one cannot discount either inertial or viscous forces. If the scaled model and original system exhibit identical *Re*, the relative importance of inertial versus viscous forces is matched between the two and any qualitative features of the flows (e.g. streamlines) will also be identical.

Dynamically scaled physical models exhibiting the same *Re* as the original systems have been used in a number of biological studies. Vogel and La Barbera (1978) outline the principles of dynamic scaling: to obtain the same *Re* when enlarging small organisms, the fluid flow must be slower and/or the fluid more viscous, and when making smaller models of large organisms, the fluid flow must be faster and/or the fluid less viscous. For instance, Vogel (1987) used air in place of water flowing at lower speeds when investigating the refilling of the squid’s mantle during swimming by scaling a model up times relative to the animal’s actual size. Vogel (1973) also investigated prairie-dog burrow ventilation using reduced scaled models and faster air flow, and more recently, Stadler et al (2016) investigated sand inhalation in skinks with 3D-printed enlarged models, using helium instead of air (thereby increasing viscosity) as the experimental fluid. Koehl and colleagues have studied crustacean antennule flicking (by lobsters (Reidenbach et al., 2008), mantis shrimp (Stacey et al., 2002) and crabs (Waldrop et al., 2015)), as well as the movements of copepod appendages (Koehl, 1995) with enlarged models, using mineral oil in place of water. The intricacies of drosophila flight have been studied using dynamically scaled robots flapping in mineral oil instead of air (Dickinson et al., 1999; Muijres et al., 2015). Finally, perhaps the largest change in scale was employed by Kim et al (2003), who modelled the bundling of *E. coli* flagella at a scale factor of ∼61,000, submerged in silicone oil (10^5^ times more viscous than water), and rotated at 0.002 rpm compared to the 100 rpm observed in real bacteria (Sowa & Berry, 2008).

In all the above studies, basic kinematics such as speeds in the original system were relatively easy to measure, and the experiments aimed to reveal the forces involved (e.g. hydrodynamic drag) or details of the fluid flow such as the pattern of streamlines. Since the representative speed *U* of the original system was known, designing experiments to achieve similitude was relatively straightforward because the *Re* was also known *a priori* – in these cases, the model size, speed, and working fluid properties were simply interrelated through *Re* (Eqn 1). For instance, once a working fluid and the model size were chosen, the required towing speed was obvious. However, in the case of sedimentation of small particles (e.g. spores, seeds, plankton), the sinking speed *U* is the key unknown. With an unknown sinking speed, the operating *Re* is also unknown, so it is not straightforward to design experiments that achieve similitude with the original system. Here we present an iterative methodology leveraging 3D printed dynamically scaled models that allows determination of the sinking speeds of small objects of arbitrarily complex shape.

We use planktonic foraminifera as an example of a small (500 – 1500 µm) biological particle for which the settling velocity is important and typically unknown. Foraminifera are a phylum of marine ameboid protists (B. K. Sen Gupta (ed.), 2002; Schiebel & Hemleben, 2005), living both on the ocean floor (benthic) and in the water column (planktonic). By secreting calcium carbonate, foraminifera produce a multi-chambered shell (test) which can grow up to 1500 µm in diameter, and which frequently exhibits complex shape. Once the organism dies or undergoes reproduction, the empty test sinks to the ocean floor, and so oceanic sediments contain substantial numbers of foraminifera tests.

Foraminifera account for 23-56% of the oceans’ production of carbonate (CO3) (Schiebel, 2002), an important factor in climate change models (Passow & Carlson, 2012). Of particular interest for climate predictions is calculating the flux of tests reaching the ocean floor (Schiebel, 2002; Jonkers & Kucera, 2015). While there are more than 30 extant species and over 600 species in the fossil record, settling velocities are known for only 14 species of foraminifera, for which *U* = 3.41×10^−4^ - 6.8×10^−2^ ms^-1^ and *Re* ≈ 18 – 55 (Fok-Pun & Komar, 1983; Takahashi & Be, 1984; Caromel et al., 2014).

## Materials and Methods

### Similitude and Settling Theory

We assume that the size (i.e., *L* – defined as the maximum length parallel to the settling direction, *A* – defined as the projected frontal area, and *V* – the particle volume not including any fluid-filled cavities), 3D shape (*Ψ*, here treated as a categorical variable due to our consideration of arbitrarily complex morphologies), and density (*ρ*_*particle*_) of the original sinking particle are known, while the sinking speed (*U*) is unknown. The properties of the fluid surrounding the original particle (i.e. *ρ*_*fluid*_, *μ*) are also known, and our goal is to design experiments in which we sink a scaled-up model particle in a working fluid of known *ρ* _*fluid*_ and *μ* in order to determine the model particle’s sedimentation speed and, via similitude, *U* of the original particle. While previous work (Berger et al., 1972; Fok-Pun & Komar, 1983; Takahashi & Be, 1984; Caromel et al., 2014) suggests that the *Re* of sinking forams should be 10^0^ – 10^2^, the exact value for morphology *Ψ* is assumed to be unknown. Hence, it is not immediately clear what size the model should be (i.e., the scale factor *S* = *L*_*model*_/*L*_*real*_) in order to match this *Re* in the experiments and ensure similitude additional constraints are required.

Throughout, we use a superscript O (“operating point”) to refer to the values of dimensioned variables at life size (e.g., 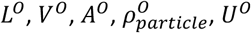) and 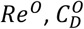 for the values of the dimensionless Reynolds number and drag coefficient (defined below) corresponding to real particles sinking in the original fluid (e.g., seawater of 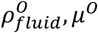). While the fluid dynamics of flow around a particle of particular shape *Ψ* can be considered theoretically over a range of *Re*, only the dynamics at *Re*^*O*^ and 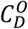 will represent the operating point corresponding to the life size particle settling speed *U*^*O*^.

When a particle is sinking steadily at its terminal velocity, the sum of the external forces acting on the particle is zero (Eqn 2); that is, the upward drag force (*F*_*drag*_, Eqn 3) and buoyant force (*F*_*buoyancy*_, Eqn 4) must balance the weight of the particle (*F*_*weight*_, Eqn 5):

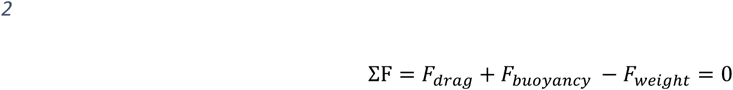

where

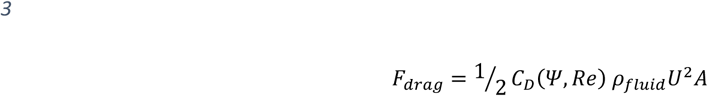

Eqn 3 introduces the drag coefficient *C*_*D*_(*Ψ, Re*), a dimensionless descriptor of how streamlined an object is. Both *C*_*D*_ and *Re* must be matched to achieve similitude. *C*_*D*_ depends on the shape of the object *Ψ*, including its orientation relative to the freestream flow – for instance, *C*_*D*_ of a flat plate oriented parallel to laminar flow is as low as 0.003 while *C*_*D*_ of a flat plate oriented perpendicular to the flow is ∼ 2.0 (Munson et al., 1994). However, in addition to object geometry, *C*_*D*_ also depends on qualitative characteristics of the flow, such as whether it is laminar or turbulent – that is, *C*_*D*_ also depends on *Re. C*_*D*_ of a sphere decreases from ∼200 at *Re* = 0.1 to ∼0.5 at *Re* = 1000; *C*_*D*_ generally decreases with *Re* for most shapes (Munson et al., 1994; Morrison, 2013). While *C*_*D*_ does not depend on object size directly, larger objects generally experience higher drag forces and this is captured by the inclusion of particle area (*A*) in the expression for *F*_*drag*_ (Eqn 3). For brevity, we will omit *Ψ* hereafter and write the drag coefficient as *C*_*D*_(*Re*).

The buoyant force (*F*_*buoyancy*_, Eqn 4) and weight (*F*_*weight*_, Eqn 5) are both expressed using particle volume (*V*_*particle*_), gravitational acceleration (*g*), and density of the fluid (*ρ*_*fluid*_) or particle (*ρ*_*particle*_), respectively:

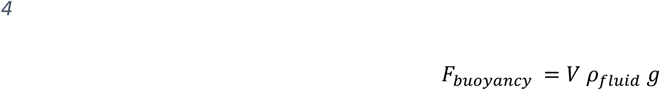

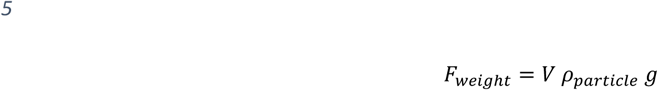

Substituting Eqns 3, 4, and 5 into 2 and eliminating *U* via the definition of *Re* (Eqn 1) yields an expression for the drag coefficient obtained through a force balance (indicated by a superscript F):

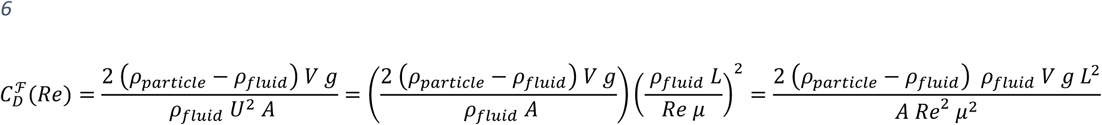

Note that this expression can be simplified further upon identification of the dimensionless Archimedes number 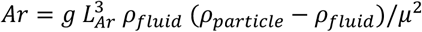 if the cubed length scale 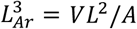, yielding 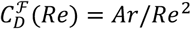, as previously highlighted by others (e.g. Karamanev, 1996). However, we will proceed with the original form of Eqn 6 to keep key variables such as *L* explicit.

If *C*_*D*_ were known for a particular foram morphology, we could simply substitute values corresponding to the original test in seawater into Eqn 6 and solve for *Re* = *Re*^*O*^ and thus *U*^*O*^ via Eqn 1, immediately solving the problem of unknown settling speed. Unfortunately, the complex shapes of foraminifera coupled with the implicit dependence of *C*_*D*_ on *Re* means that both variables are generally unknown, and thus far we have only one constraining relationship between 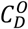 and *Re*. More information is required to determine where along this constraint curve the operating *C*^*O*^ and *Re*^*O*^ are located. This information can come from experiments in which the sinking speeds of scaled-up model particles of various sizes (i.e., scale factors *S*) in a viscous fluid are measured directly, allowing us to calculate *Re* via Eqn 1 and then 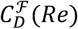 via Eqn 6 for the models, with appropriate values substituted for each experiment. For clarity, we can rewrite Eqn 6 for a model in terms of *S* and the original test parameters (*L*^*O*^, *A*^*O*^, *V*^*O*^):

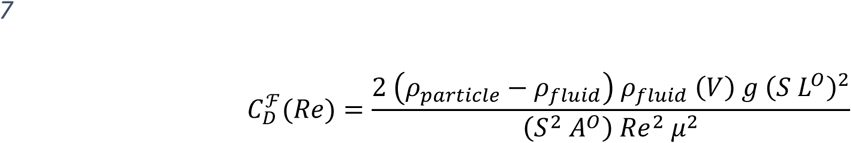

where we use the fact that for a model, *L* = *S L*^*O*^ and *A* = *S*^2^*A*^*O*^. While one would also expect *V* = *S*^3^ *V*^*O*^ for 3D printed models, limitations of our 3D printer led to variation in *V* that we overcame using a more general empirical relationship between *S* and *V* based on mass measurements – see 3D Printer Limitations. Eqn 7 represents a constraining relationship between *C*_*D*_ and *Re* for the sinking particle, which we use to collect (*Re, C*_*D*_) experimental data points at several *S*. Once sufficient data are collected, we can construct a new, empirical relationship (e.g., a cubic spline fit) between *C*_*D*_ and *Re* for a particular particle shape, which we term 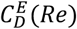. Finally, we can solve for the operating *Re*^*O*^, 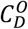, and *U*^*O*^ by finding the intersection point between the 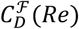 constraint curve specific to life-size particles sinking in seawater (i.e., Eqn 7 with *S* = 1 and 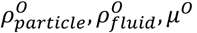) and our empirical 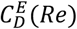 spline curve valid for a particular particle shape moving steadily through any fluid.

### Overview of experiments

To construct an empirical 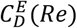 curve for a particular test morphology, we started with 3D scans of individual specimens (SI, Table 1). These data allowed us to easily fabricate scaled-up (scale factor *S*) physical models of each specimen using a desktop 3D printer (SI 2). The models were then released in a tank of mineral oil (*ρ* = 830 kg m^-3^, *μ* = 0.022 Pa S) and imaged by two digital cameras oriented at 90 degrees to each other to record 3D trajectories (SI 3, SI Fig. 7). There were two primary difficulties with these experiments: the first associated with the resolution of our 3D printer, and the second associated with wall effects, which were overcome as follows.

### 3D Printer Limitations

Whilst in principle, the volume of a printed model should simply scale according to *V* = *S*^3^*V*^*O*^, due to inherent limitations of the 3D printer as well as difficulty in removing excess resin from small models, we found that this expectation was usually not satisfied, and weighing the models showed that *M*/*ρ*_*particle*_ > *S*^3^ *V*^*O*^ (SI Fig. 3). Therefore, we estimated *V* of each model by weighing on an Entris 224-1S mass balance (±0.001 g) and assuming *ρ*_*particle*_ was 1121.43 ± 13.73 kg m^-3^, based on the average mass of five 1 cm^3^ cubes of printed resin. Furthermore, whenever a predicted value for *V* at a given scale factor *S* was needed, i.e. in Eqn 7 (see Remaining iterations), we based this on cubic spline interpolation of our *V*(*S*) data for existing models when sufficient data were available, with extrapolation based on cubic scaling of *V*(*S*) if required (see SI Fig. 3):

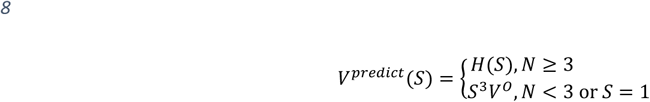

where *N* is the number of existing volume measurements and *H* represents the cubic spline fit of *V* vs *S*. Note that because we always directly measured *V* by weighing after printing each model, and it is not necessary to achieve the exact *Re* and *C*_*D*_ of the operating point (*Re*^*O*^and 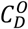) in the experiments (see Remaining iterations), this correction was not strictly required for our method to succeed. It merely aids in improving the rate of convergence of our iterative approach by reducing the difference between our anticipated and actual *Re, C*_*D*_ for each experiment.

### Wall effects

At low *Re*, the effects of artificial walls in an experimental (or computational) system can be nonintuitively large and lead to substantial errors if not accounted for (Vogel, 1994). Acting as an additional source of drag, the walls several tens of particle diameters away can slow down a sinking particle and increase its apparent drag coefficient. We designed our experiments to minimise wall effects by using an 0.8 m diameter tank (SI Fig. 6) and model diameters on the order of 1 cm. To reduce errors further, we applied the method of Happel and Brenner (1983) to convert between the apparent drag coefficient when walls are present 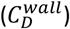 and the desired drag coefficient in an unbounded domain 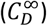:

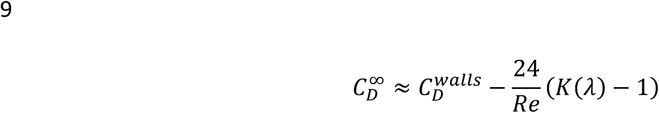

where

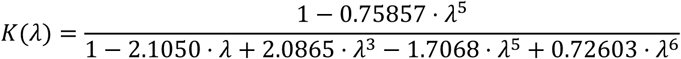

Here, *λ* = *d*/*D* where *d* is the diameter of the sinking particle and *D* is the tank diameter; we take *d* = *L*. While Eqn 9 is not exact, it substantially reduces the error otherwise incurred if one were to neglect wall effects entirely. We applied this correction by taking any experimentally determined *C*_*D*_ to equal 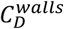, and using 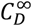 estimated according to Eqn 9 for subsequent calculations as detailed below.

### Iterative approach

#### First iteration

To construct an empirical cubic spline 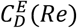 needed to solve for *U*^*O*^, at least three experimental data points (corresponding to three scale factors) are needed. These first three *S* were chosen by using an existing empirical *C*_*D*_(*Re*) relationship for a sphere, valid for 0 < *Re* < 10^6^ (Morrison, 2013, Fig. 8.13, page 625):

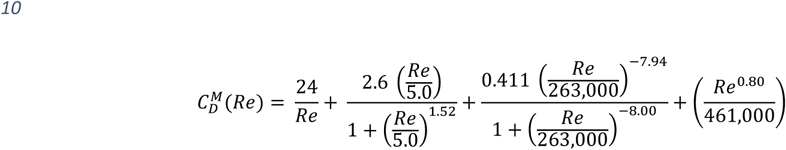

While morphologically complex particles such as foraminifera tests are not expected to behave like ideal spheres, Eqn 10 should be sufficient to provide initial guesses, after which we iterate to find the solution.

Substituting Eqn 10 into Eqn 7 (with *S* = 1, *V* = *V*^*O*^, and 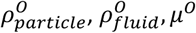 substituted) and moving all terms to one side, we can numerically solve (MATLAB: *fzero* function) for our first estimate of the operating *Re*^*O*^. Substituting this *Re* back into Eqn 7 or Eqn 10 yields an estimate of the operating 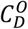. We aimed to reproduce this *Re* and *C*_*D*_ in the first experiment, excepting that we accounted for wall effects by distinguishing between 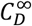 and 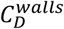 expected to occur in the tank. Hence, we could again substitute this *Re* into Eqn 7 but now with *ρ*_*particle*_ corresponding to the resin model and *ρ*_*fluid*_ and *μ* corresponding to mineral oil, and combine this expression with Eqn 9, assuming our estimated 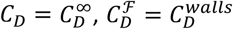, and *λ* = *S L*^*O*^ /*D*. The resulting expression can be solved numerically for the first scale factor, termed *S*_1_. Two more scale factors (*S*_2_ and *S*_3_), one smaller and one larger than *S*_1_, were chosen to span expected *Re* values for forams from published literature (e.g. Fok-Pun & Komar, 1983; Takahashi & Be, 1984; Caromel et al., 2014) as well as *Re*^*O*^ for other species which had reached convergence. This procedure was intended to bound the correct *S* value that reproduces the operating *Re*^*O*^ and 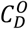 of the settling particle. The three models were printed, their actual volumes *V* measured via weighing, and their settling velocities *U* experimentally measured as detailed in SI 3.

An empirical cubic spline curve 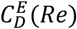 can now be fitted (D’Errico, 2009) to these three initial (*Re, C*_*D*_) data points, constrained to be monotonically decreasing and concave up within the limits of the data to match expectations for drag on objects at low to moderate *Re*. Three optimally spaced spline knots were used since this yielded excellent fits to the data as the number of data points increased. These details of the spline as well as its order (i.e., cubic vs linear) are somewhat arbitrary but we ensured that our results were sufficiently converged as to be insensitive to them (see Remaining iterations).

The operating point 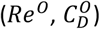 corresponding to the particle settling in the natural environment can be visually represented as the intersection point of the 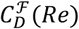 curve defined by Eqn 7 (with *S* = 1 and 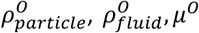) and the empirical 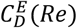 relationship based on our experimental data. Algebraically, the operating point is the solution to 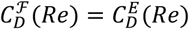. We solved for *Re*^*O*^ numerically using a root finding algorithm (MATLAB’s *fzero*) on the objective function 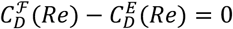 and then obtained 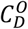 by substituting *Re*^*O*^ into Eqn 7. Finally, *U*^*O*^ was easily determined from the definition of *Re*^*O*^ (Eqn 1 with *U*^*O*^, *L*^*O*^, and 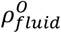 substituted).

Since our first three empirical data points and fitted spline 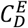 corresponded to guessed model scale factors *S*, our initial operating point prediction 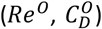 often was not located near any of these initial points or sometimes even within the bounds of these data (in which case linear extrapolation of 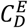 was used to estimate the operating point). Therefore, to ensure the accuracy of our predicted *U*^*O*^, we continued iterating with additional experiments.

#### Remaining iterations

The model scale factor for the *N*^th^ experiment was chosen by combining Eqns 7, 8, and 9 with *Re* = *Re*^*O*^ and 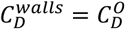, (from the previous iteration), 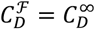, and *V* = *V*^*predict*^, and numerically solving for *S*. A model close to this new scale was printed and sunk, its settling velocity *U* recorded and *Re* and *C*_*D*_ computed, and a more accurate spline 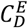 constructed by including this new data point. The calculation of 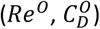 detailed in the previous section was then repeated, yielding a more accurate operating point. Overall, the aim was to tightly bound the predicted operating point with experimental data to maximize confidence in the fitted spline in this region.

1. The iterative process (visualized as a flowchart, Fig. 1) was repeated until:
2. the predicted operating point was not extrapolated beyond our existing data,
3. the variation in calculated *U*^*O*^ between the fitting of a linear spline and cubic spline was no greater than 5%, and
4. the variation between the predicted *Re*^*O*^ and the closest experimentally measured *Re* was less than 15%.

Through this method we calculated the sinking velocities of 30 species of planktonic foraminifera (Table 1).

**Table 1.** Predicted sinking speeds U^O^ for the 30 species of planktonic foraminifera included in this study. 1 indicates species scanned at PETRA III synchrotron storage ring at DESY in Germany. Scans of the remaining 27 species were obtained from The University of Tohoku museum’s database. Operating Re^O^ and 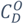 predicted for each species are also presented, and for comparison, the theoretical 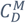 of a sphere at the same Re^O^. The number of model iterations required to achieve convergence of U^O^is listed. Species morphologies are shown in SI Fig. 2.

| Species | $L^0$<br>($\mu\text{m}$) | $U_{particle}^0$<br>($\text{cm s}^{-1}$) | $Re^0$ | $C_D^0$ | $C_D^M(Re^0)$ | Number<br>of<br>iterations |
| --- | --- | --- | --- | --- | --- | --- |
| *Neogloboquadrina acostaensis* | 307 | 3.6 | 9.3 | 2.5 | 4.16 | 4 |
| *Globigerinella adamsi* | 596 | 3 | 9.6 | 2.6 | 4.08 | 4 |
| *Dentoglobigerina altispira* | 282 | 6.1 | 37.3 | 1.6 | 1.76 | 5 |
| *Globoturborotalita apertura* | 650 | 2.8 | 7.4 | 4.2 | 4.82 | 3 |
| *Globigerina bulloides* | 296 | 3.3 | 13.4 | 2.4 | 3.29 | 5 |
| *Globigerinoides conglobatus*<sup>1</sup> | 445 | 1.8 | 5.8 | 5.9 | 5.67 | 4 |
| *Truncorotalia crassaformis* | 250 | 3.3 | 9.5 | 2.3 | 4.09 | 4 |
| *Praeorbulina curva* | 433 | 4.8 | 17.8 | 2.2 | 2.76 | 3 |
| *Sphaeroidinella dehiscens* | 460 | 5.1 | 32.8 | 1.7 | 1.9 | 4 |
| *Sphaeroidinella dehiscens*<sup>1</sup> | 269 | 2.5 | 8.3 | 5 | 4.48 | 4 |
| *Catapsydrax dissimilis* | 380 | 6.2 | 36.4 | 1.7 | 1.78 | 4 |
| *Neogloboquadrina dutertrei* | 259 | 4 | 17.6 | 2 | 2.77 | 3 |
| *Globigerinoides fistulosus* | 392 | 2.6 | 11.2 | 3 | 3.7 | 4 |
| *Globorotaloides hexagonus* | 321 | 2.7 | 8.3 | 3.4 | 4.48 | 4 |
| *Neogloboquadrina humerosa* | 360 | 5 | 22.7 | 2 | 2.37 | 4 |
| *Globoconella inflata* | 422 | 3.9 | 21.4 | 2.5 | 2.45 | 4 |
| *Fohsella lobata* | 312 | 4.2 | 13.1 | 1.5 | 3.34 | 3 |
| *Hirsutella margaritae* | 258 | 2.9 | 8.1 | 2.8 | 4.55 | 3 |
| *Paragloborotalia mayeri* | 298 | 3.1 | 8.1 | 3.2 | 4.54 | 4 |
| *Globoturborotalita nepenthes* | 353 | 4.2 | 16.4 | 2.4 | 2.9 | 3 |
| *Pulleniatina obliquiloculata* | 350 | 4.1 | 24.6 | 2.1 | 2.26 | 3 |
| *Fohsella peripheroronda* | 422 | 3.6 | 10.1 | 2.4 | 3.95 | 3 |
| *Globigerina praebulloides* | 539 | 2.8 | 9.4 | 3.2 | 4.14 | 4 |
| *Menardella praemenardii* | 569 | 3.9 | 14 | 2 | 3.2 | 4 |
| *Globoconella puncticulata* | 269 | 4 | 14.5 | 2.8 | 3.14 | 3 |
| *Fohsella robusta* | 312 | 3.5 | 11.2 | 2.3 | 3.69 | 4 |
| *Dentoglobigerina rohri* | 303 | 5.7 | 33.6 | 2 | 1.87 | 3 |
| *Paragloborotalia siakensis* | 366 | 3.4 | 10.4 | 2.6 | 3.86 | 4 |
| *Globoconella sphericomiozea* | 627 | 4.3 | 18.8 | 2.3 | 2.66 | 4 |
| *Truncorotalia truncatulinoides* | 325 | 6.3 | 42.4 | 1.5 | 1.63 | 6 |
| *Orbulina universa*<sup>1</sup> | 567 | 1 | 3.3 | 10.7 | 8.67 | 6 |

**Fig. 1.**
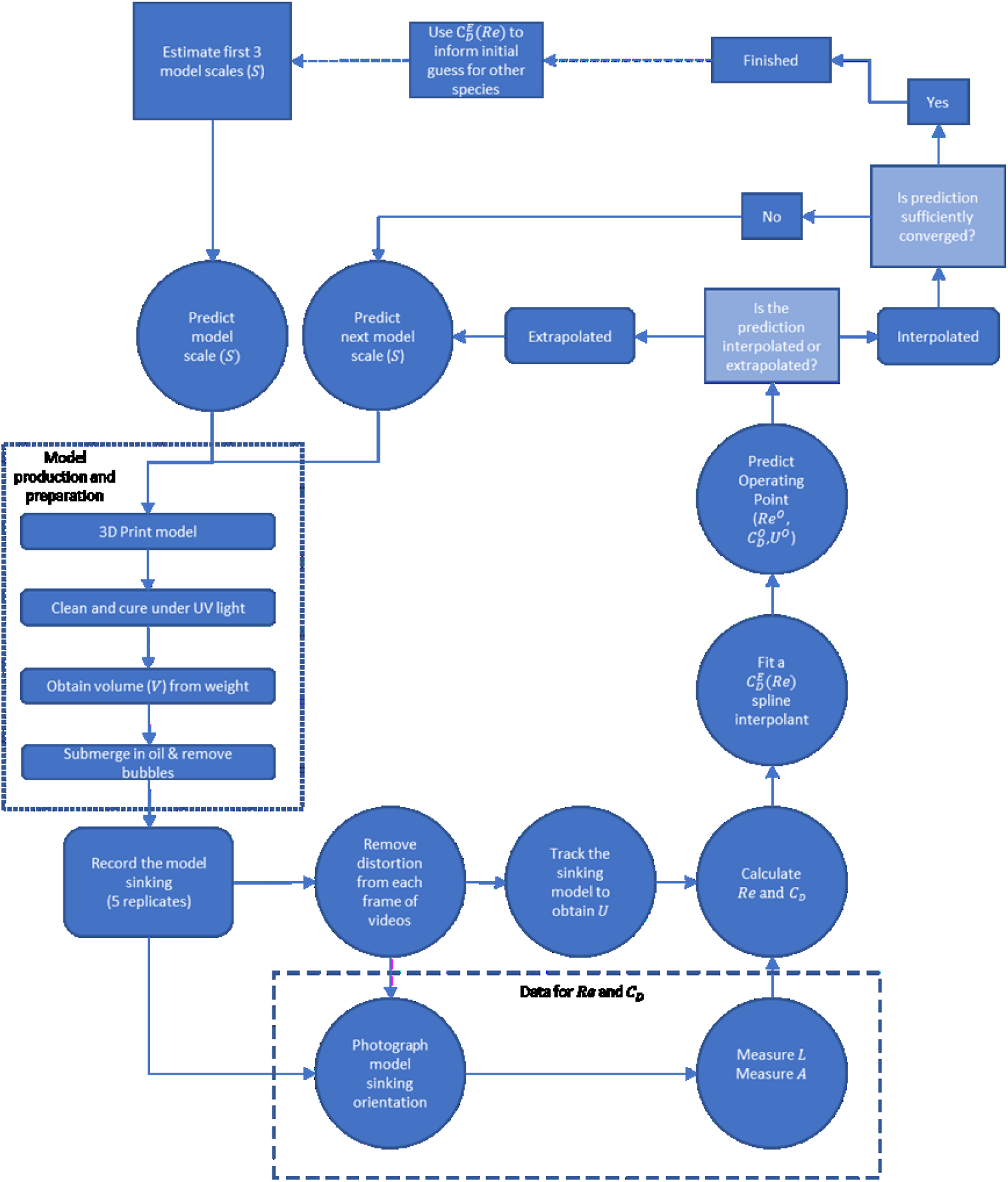
Summary of the full method; details are discussed in main text. Lighter shades represent a decision, square boxes are data inputs, rounded square boxes are manual processes, and circles are computational steps.

## Results

Our basic methodology was first validated by 3D printing a series of spherical models (10 – 20 mm in diameter) for which the theoretical *C*_*D*_(*Re*) relationship is already well-known. In order to achieve low density (and thus low sinking velocity and low *Re*), these spheres were hollow and filled with oil via two small holes (of diameter 0.8% of the sphere diameter). Our empirically generated 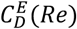 curve compares favourably with the theoretical 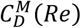 curve (Morrison, 2010) (R^2^=0.875, SI Fig. 5), with the distance between the curves approximately constant above *Re* ≈ 25. While the error grows larger at lower *Re*, we expect most foram species to operate at *Re* ≈ 18 – 55 based on previous work (Berger et al., 1972; Fok-Pun & Komar, 1983; Takahashi & Be, 1984; Caromel et al., 2014).

To quantify errors in our approach even more directly, we then considered hypothetical hollow spherical particles with the same material density 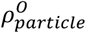 as foram tests and a range of sizes (*L*^*O*^ = 750 − 1150 μm, similar to the species we studied) settling in seawater. This size range corresponds to *Re* = 12 − 27, the area where our 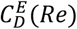 curve is most divergent from 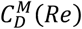. We compared predictions of the operating *U*^*O*^ based on our empirical 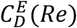 curve versus the theoretical 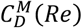 curve for spheres as outlined above, substituting Eqn 10 for 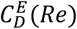 in the latter case. Maximum relative error in predicted *U*^*O*^ was 11.5% at *Re* = 16 (corresponding to a sphere 860 µm in diameter) while the minimum difference was 6.5% at *Re* = 27 (corresponding to a sphere of 1150 µm in diameter, SI Fig. 5). This level of error is much smaller than the variation in *U*^*O*^ we predicted across the 30 foram species we investigated (Table 1).

There was little variation in the number of iterations required to reach convergence (mean 4, range 3-6, see Table 1), despite the morphological complexity of some species (e.g. *Globigerinoides fistulosus*). We suspect the higher end of this range was due to these species having forms that were particularly challenging to clean residual resin from, or the incomplete removal of air bubbles once submerged in oil. In Fig. 4 we present an example of convergence of our method to the operating *Re*^*O*^, 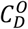, and *U*^*O*^. Comparisons with previously published data suggest that our values lie within the expected range of sinking foraminifera (Berger et al., 1972; Fok-Pun & Komar, 1983; Takahashi & Be, 1984; Caromel et al., 2014, SI Fig. 1).

## Discussion

Here we present a novel method of determining settling speed by leveraging dynamically scaled models falling under gravity rather than being towed at a controlled speed. Applying our method to foraminifera-inspired spherical particles, we predict settling speeds within 11.5% of theoretical expectations (SI Fig. 5). Our predicted speeds of real foraminifera also fall within existing data for 14 species (SI Fig. 1, Fok-Pun & Komar, 1983; Takahashi & Be, 1984; Caromel et al., 2014) and compare well with known sinking speeds for other particles of comparable size and density (e.g. faecal pellets, (Table 3, Iversen & Ploug, 2010), various phytoplankton (Fig. 1, Smayda, 1971)).

Key questions in biological oceanography revolve around the sedimentation of both living and dead planktonic organisms. Sedimentation of microscale plankton has been measured both *in situ* (e.g. Waniek, Koeve and Prien, 2000) and in the laboratory (e.g. Smayda, 1971; Miklasz and Denny, 2010). By settling dense suspensions of microorganisms, these studies provided a population sinking rate (Bienfang, 1981) which could be two to three times lower than the settling velocity of an isolated particle in the typically dilute ocean (Miklasz & Denny 2010). Furthermore, small particles tend to be drawn towards the walls of an experimental container (Happel & Brenner, 1983) where they experience artificially higher drag (i.e. wall effects). We overcame such issues by settling individual scaled up models in a large tank at least 45 times larger than the models, with an additional mathematical correction for wall effects.

Some previous studies have also used enlarged models of microscale plankton to facilitate observations. Padisak et al. (2003) used handmade models of plankton to examine form resistance, or how deviation in shape from a sphere of equal volume affects sinking velocity, but there was no attempt at accurately matching *Re*. Holland (2010) used mechanical pencil leads sedimenting in castor oil to examine the effect of orientation on sinking diatom chains. In an improvement compared to Padisak et al. (2003), the *Re* of the pencil lead was shown to be less than 1, as appears to be the case for sinking diatoms (Miklasz & Denny 2010). However, neither Padisak et al. (2003) nor Holland (2010) calculated sinking velocities for the real organisms. Our dynamic scaling approach ensures that we accurately recreate the fluid flows around settling organisms – a requirement for the correct prediction of sinking speeds. We also improve on these previous methodologies by obtaining volume measurements from µCT scans and by using consumer-grade cameras to observe natural sinking orientation and obtain consistent measurements of frontal surface area relative to the flow.

By design, our dynamic scaling approach yields an interpolated *C*_*D*_(*Re*) curve that describes the flow dynamics (and thus sinking speeds) that would occur if various fluid and/or particle parameters were varied, offering a degree of flexibility not seen in other studies. For example, phytoplankton blooms can increase both the density and viscosity of water due to exudates (Jenkinson et al., 2015), while increasing global temperatures have the opposite effect. The density and viscosity of seawater also naturally vary with latitude. Understanding how these variations affect sinking rates can offer insights into the evolutionary pressures on plankton.

Our method can easily be modified to study sedimenting particles operating at any *Re*, providing the system’s *Re* range is experimentally replicated. Other sinking marine particles include diatoms (*Re* ≈ 10^−2^ - 1,Botte, Ribera D’Alcalà and Montresor, 2013) and radiolaria (*Re* ≈ 10 - 200,Takahashi and Honjo, 1983), for which one could use digital models as we have in conjunction with a suitably viscous fluid (SI Table 1) to enable sufficiently large models to be produced (SI Table 2). The method can also be applied to terrestrial systems such as settling spores (*Re* ≈ 50 e.g. Noblin, Yang and Dumais, 2009) and dispersing seeds (*Re* ≈ 10^3^ Azuma and Yasuda, 1989), again by using 3D printed models based on (often existing) µCT data.

Whilst our method pertains to settling in a quiescent fluid, it would be relatively simple to conduct similar experiments using a flume to calculate threshold resuspension velocities (i.e. the horizontal flow speed required to lift a particle off the substrate), important in the study of wind erosion and particle transport and deposition (Bloesch, 1995; Bagnold, 1971). Similarly, studying particles suspended in shear flow could be achieved using a treadmill-like device as used by Durham et al (2009) or a Taylor-Couette apparatus as per Karp-Boss & Jumars (1998). While additional dimensionless groups beyond *Re* and *C*_*D*_ would need to be matched to achieve similitude in these systems, we hope that our study provides a starting point for the experimental study of these and other more complex problems.

## Supplementary Information

### SI 1. Study Species

To expand the limited number of planktonic foraminifera species for which estimates of sinking velocities exist, the 30 species used are listed in Table 1. The majority of the species were selected from the University of Tohoku museum’s database, eforam Stock (http://webdb2.museum.tohoku.ac.jp/e-foram/), with a micro computed tomography (µCT) scan resolution between 2.5 and 3.6 pixels per µm, and were exported as 3D triangular mesh (STL format) files. For species where more than one scan was available, the scan that contained the best-preserved specimen was chosen. By only including one specimen per species, this approach neglects phenotypic plasticity which is demonstrated in planktonic foraminifera (e.g. Lohmann, 1983; Morard *et al*., 2013), but was chosen due to limitations of µCT scan availability and time constraints on the project. An additional 3 species were scanned at the PETRA III synchrotron storage ring at DESY in Germany, with a scan resolution of 0.721 µm per pixel. These scans were segmented and rendered using SPIERS (Sutton et al., 2012), and again exported in STL format. Meshes of all foraminifera were manually checked in Meshlab (Callieri et al., 2012) for integrity.

**SI Fig. 1.**
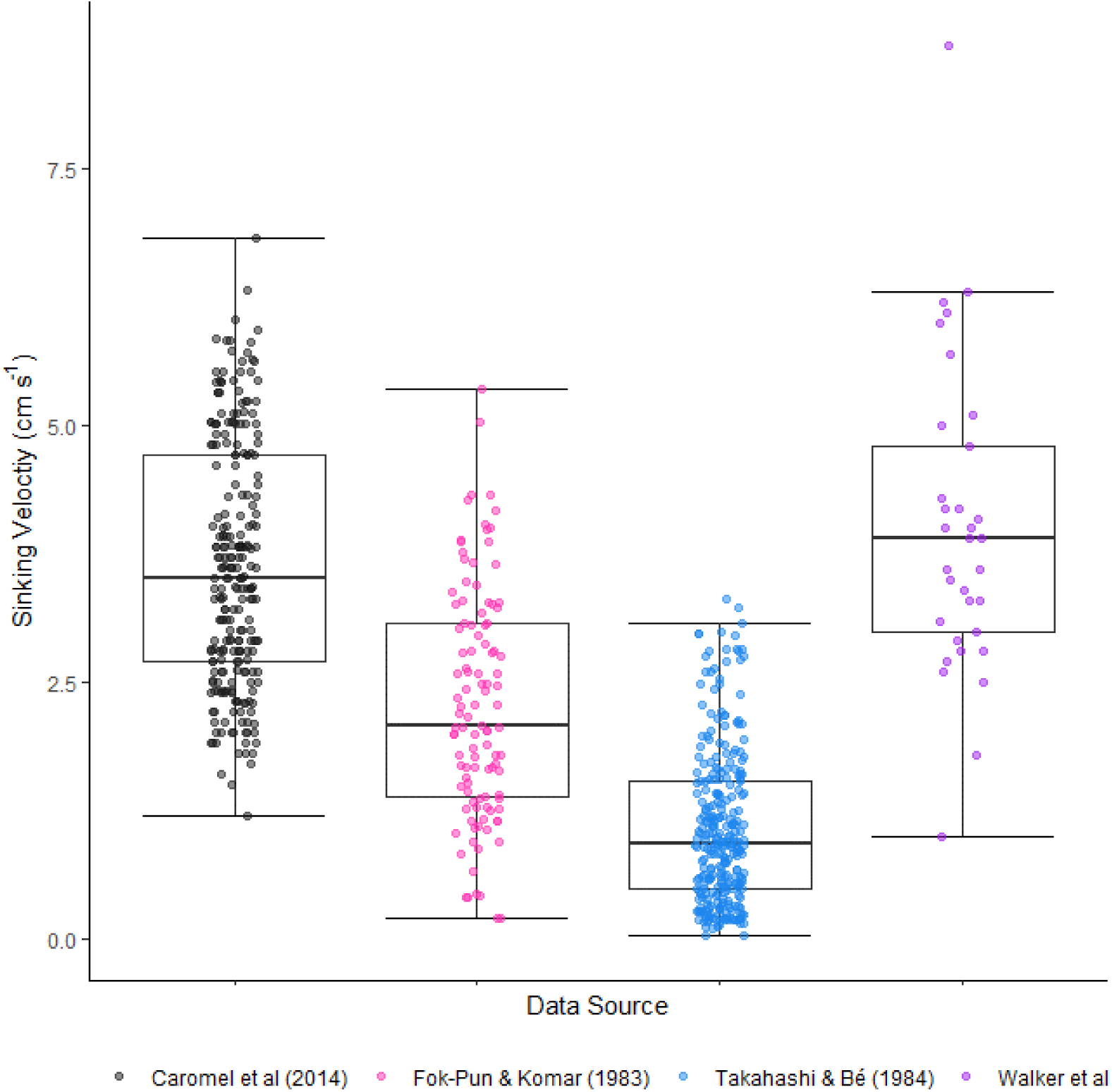
Comparison of sinking velocities between other studies (Caromel et al., 2014; Takahashi & Be, 1984; Fok-Pun & Komar, 1983) and the data from this study (Walker et al). Each point represents an individual foraminifera (or model).

**SI Fig. 2:**
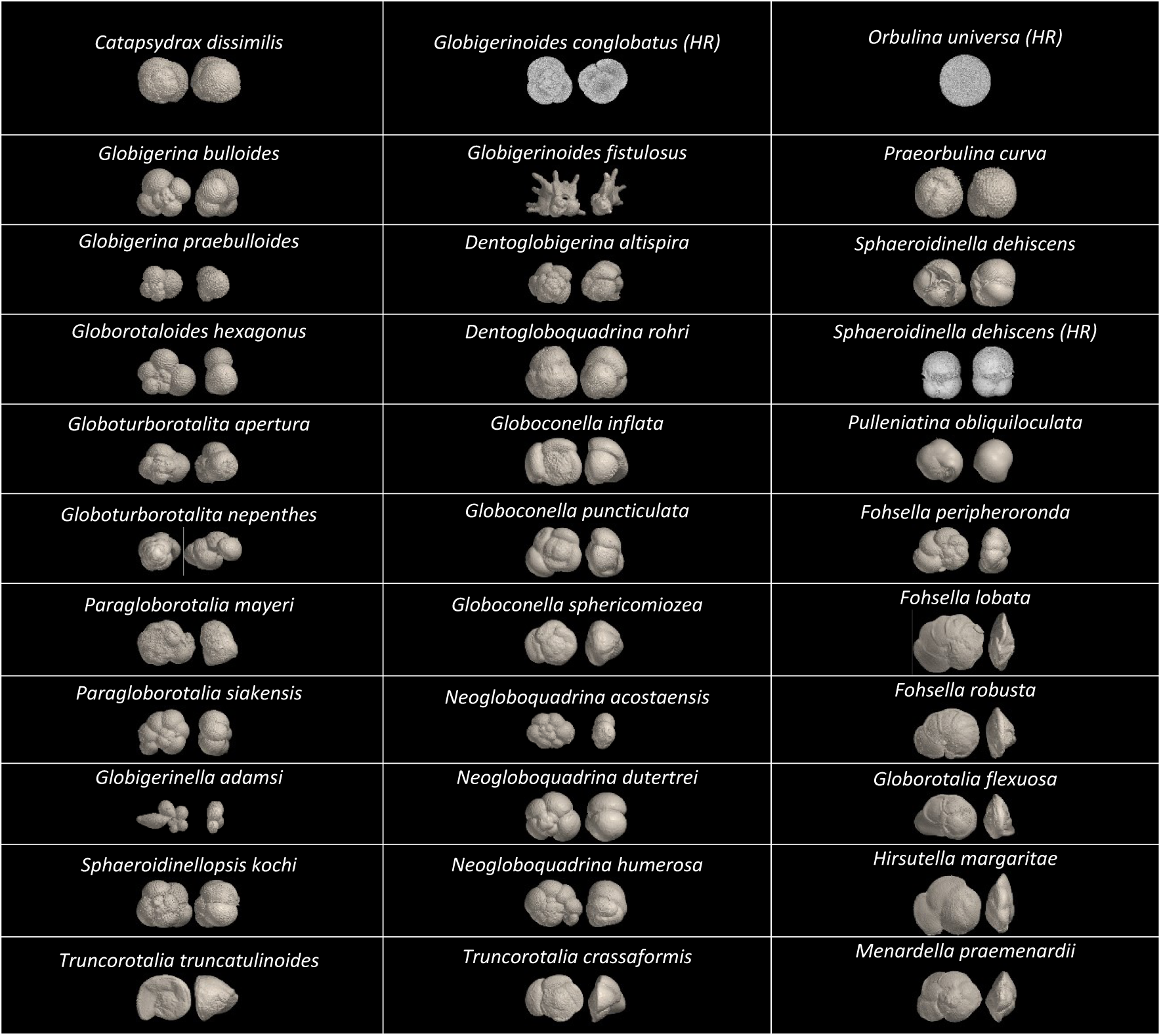
Each species studied, shown in both spiral view and 90° rotation (so that the spiral view is facing to the left of the page), note O.universa is spherical in appearance so only one view is provided. (HR) denotes species scanned at high resolution. Species are grouped according to Aze et al’s (2010) convention.

### SI 2. 3D printing and model preparation

All models were 3D printed using a FormLabs Form1+ (Formlabs, Somerville, Massachusetts, USA) 3D printer, using FormLabs Clear Resin Version 2 (Formlabs, Somerville, Massachusetts, USA) with a layer thickness of 50 µm. Models were washed and flushed with isopropanol to remove excess resin following Formlabs’ guide and allowed to air dry. Support material was removed, and the models lightly sanded with 400 grit Wet ‘n’ Dry paper, followed by a final isopropanol wash to remove any remaining residue. Once dry, models were filled with mineral oil in preparation for sinking. Clear resin was chosen to allow each model to be checked for bubbles (which would increase the buoyancy of the model). Any bubbles were removed using a 30-gauge needle and syringe.

Following convention when defining the area *A*_*particle*_ used in the definition of *C*_*D*_ (Eqn **3**), we measured projected area of the sinking foraminifera. Referring to high resolution images of the sinking model (SI Fig. 4), a digital model of the foraminifera was manually aligned to measure the projected area in a plane perpendicular to the sinking direction (Fig. [2]b). We used the same procedure to measure the maximum length parallel to the flow (*L*) for the calculation of *Re* (Fig. 2). These choices facilitated objective comparisons of *C*_*D*_ across morphologically diverse species, to be detailed in a future study.

**Fig. 2:**
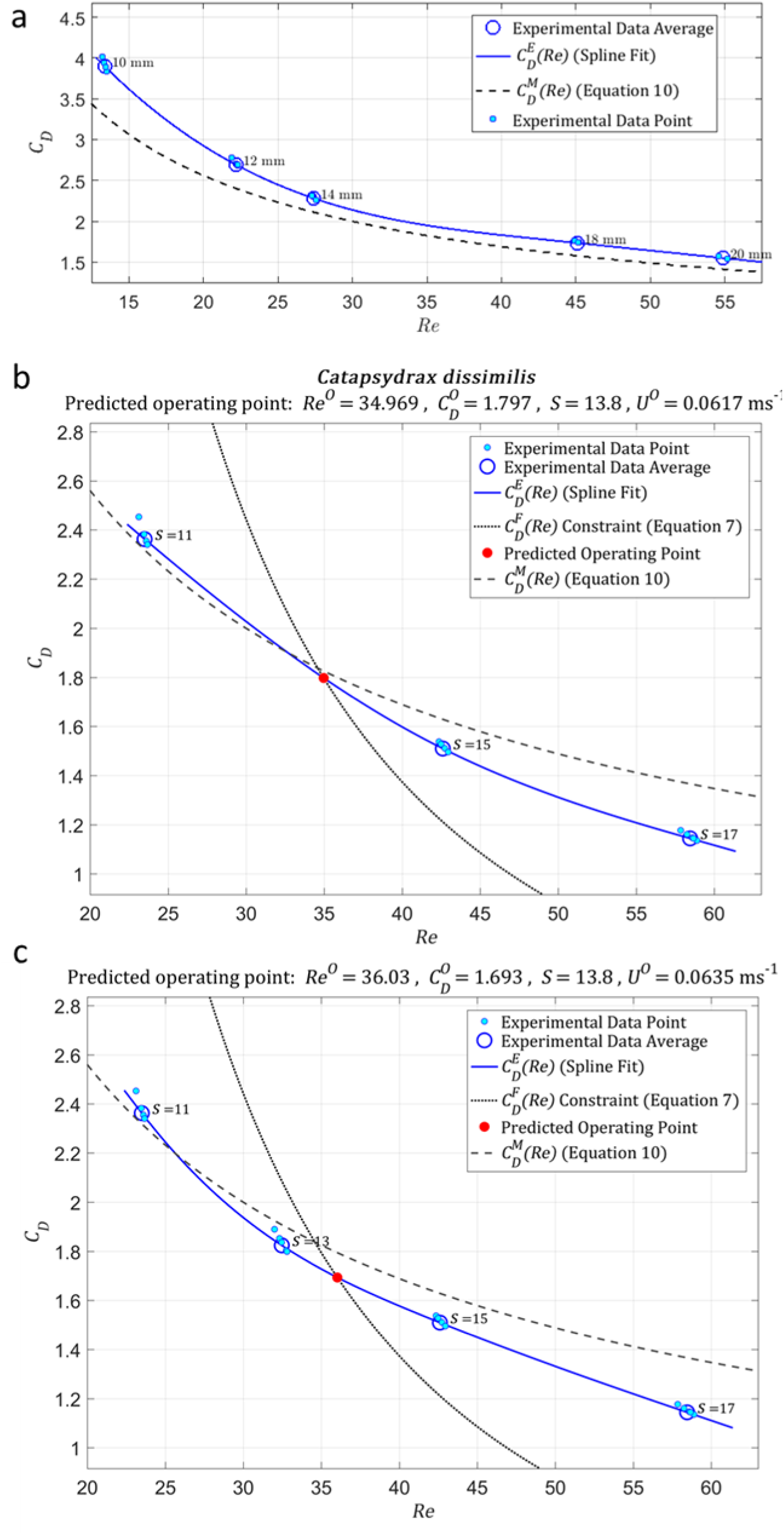
A) Comparison of our empirically generated 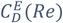 curve for 3D printed spheres versus the theoretical 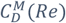 curve (Morrison, 2010). Goodness of fit of 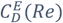 to 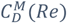 is R^2^ = 0.857. Sphere diameter is indicated for each model. Fig2. B & C) An example of our iterative solution process for Catapsydrax dissimilis (as in SI Fig. 3 showing predicted operating values (including best estimates of the required model scale S to achieve similitude) based on experimental data from 3 (A), and 4 (B) models. S for each model are labelled. The operating point is computed as the intersection where 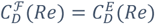. For reference, the theoretical 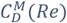 curve (Morrison, 2010) for a sphere is also shown.

**SI Fig. 3:**
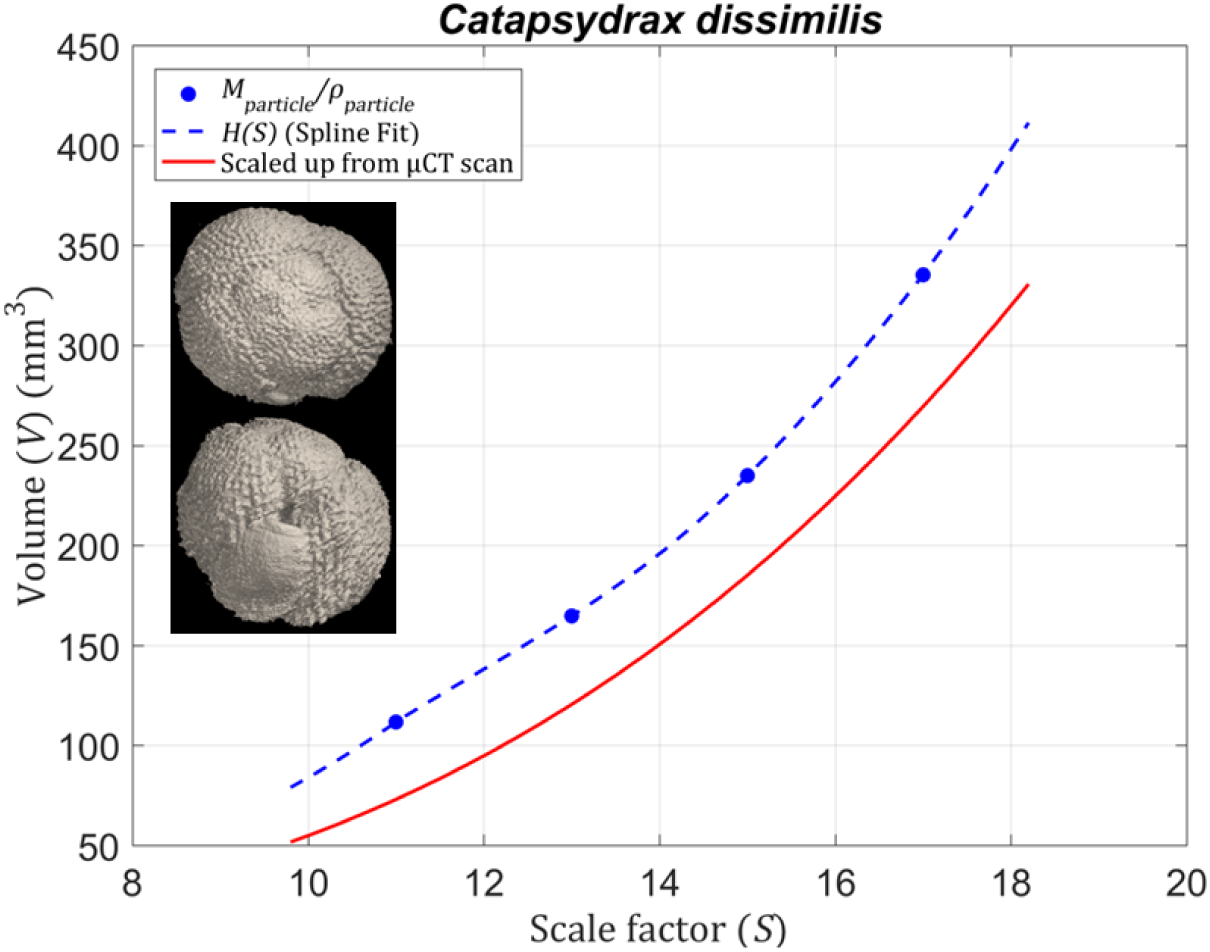
Differences in measured (via mass balance, blue dots and spline fit) and expected (scaled up from digital surface mesh, red) model volumes versus scale factor for C. dissimilis. Inset: Scanned C. dissimilis specimen from a spiral (top) and aperture (bottom) view; maximum length is 610 µm.

**SI Fig. 4:**
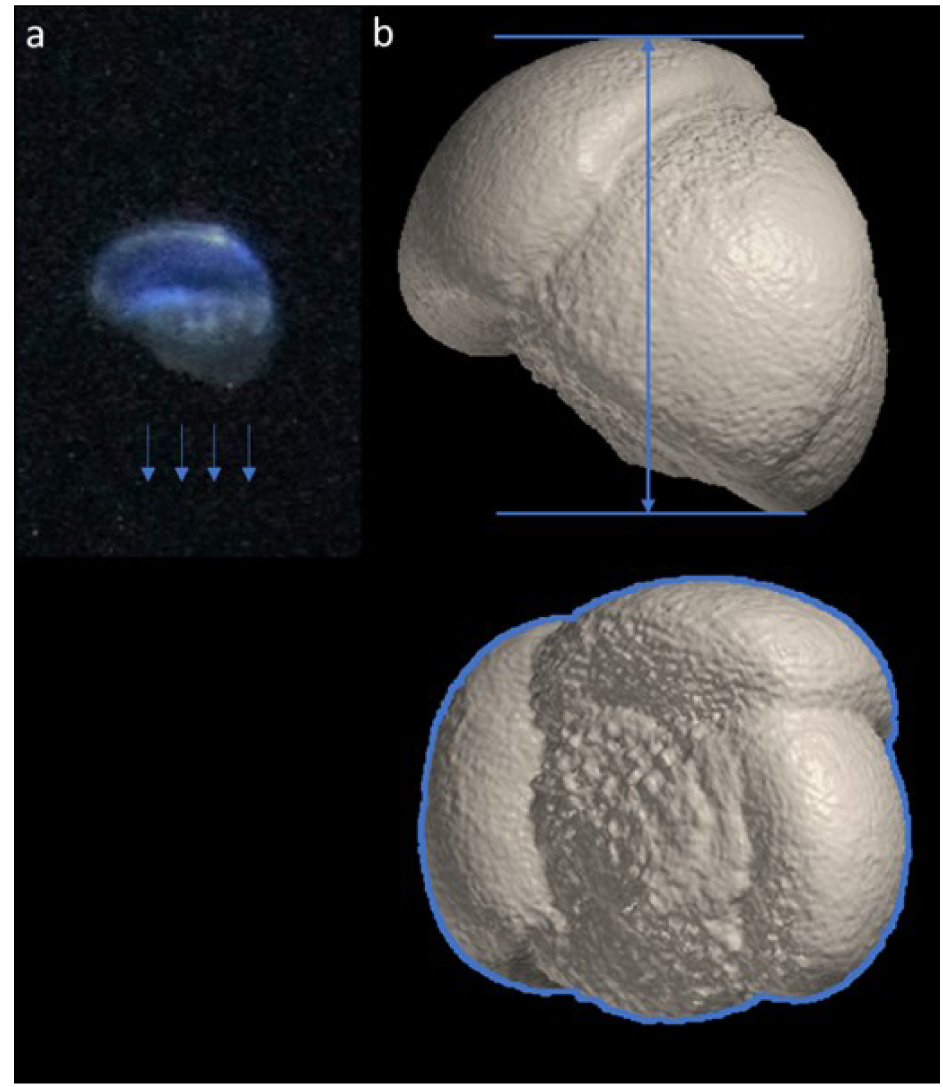
a) An example high resolution image of a sinking foraminifera model (Globoconella inflata). b) Top: The computer-generated model of the same specimen rotated to the sinking orientation, with the maximum length parallel to the flow (L). Bottom: underside view of the foraminifera, i.e. the projected area perpendicular to the flow as the foraminifera sinks, with the projected area (A) outlined in blue.

**SI Fig. 5.**
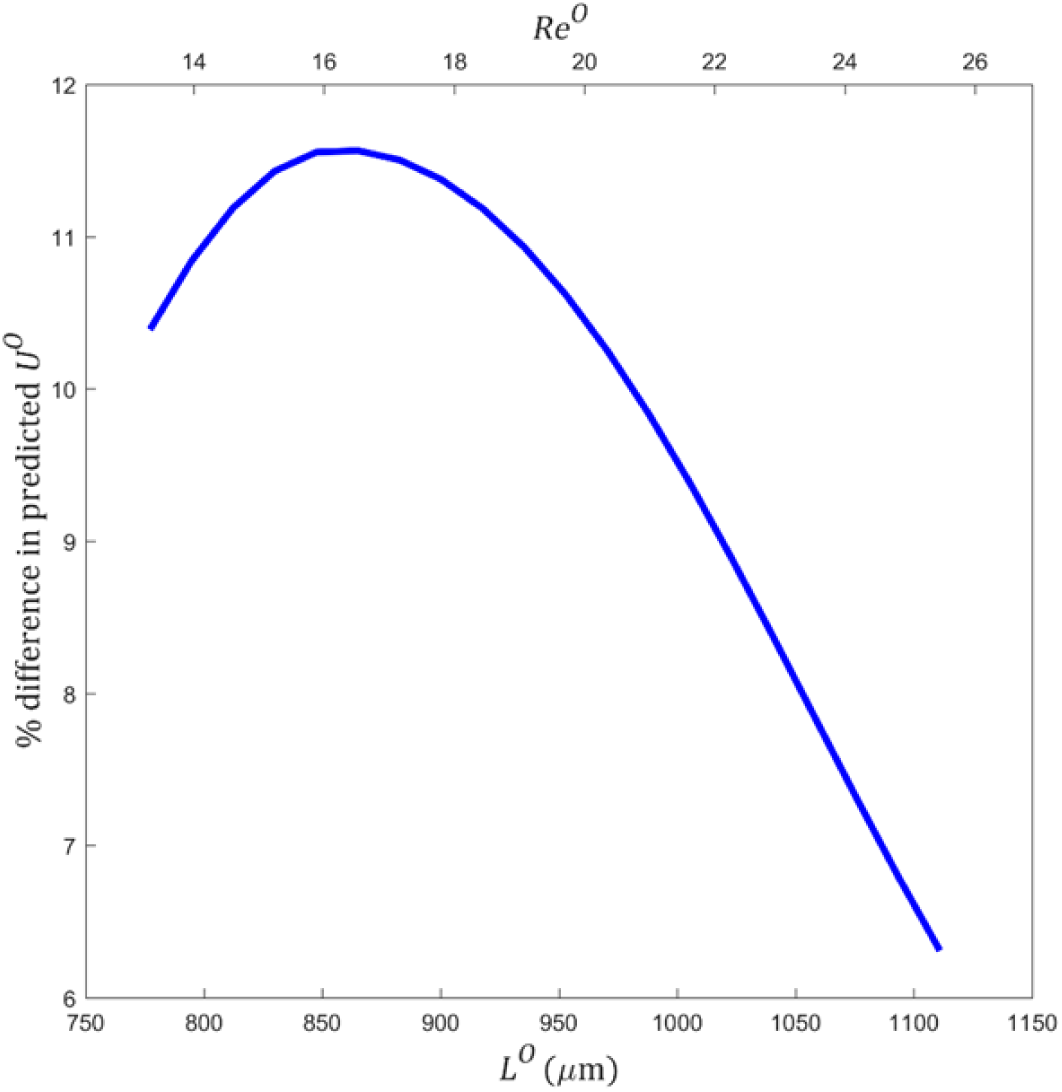
Relative difference between predicted U^O^ based on our empirical 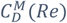) curve versus the theoretical 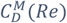 curve for hypothetical hollow spherical particles in seawater. Re^O^ is based on operating point prediction using 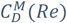.

### SI 3. Settling Tank, particle imaging, and velocity calculation

A Perspex tank 0.9 m in diameter and 1.2 m in height was filled with “Carnation” white mineral oil (Tennants Distribution Limited, Cheetham, Manchester, UK) to a depth of 1.18m (approximately 750 L). The tank was fitted with a custom net and net retrieval system (SI Fig. 6) to allow easy retrieval of the models after their descent, allowing each model to be sunk 5 times. Integrated to the net retrieval system was the release mechanism, which was held centrally over the tank, with the grasping parts submerged below the oil level. This ensured that the model was released in a controlled and repeatable fashion.

To minimise reflections, the tank was surrounded by a black fabric tent-like structure. This also served as a dark background to facilitate visualisation of the model during descent. The tank was illuminated with a single 800 lumen LED spotlight was placed underneath the tank and, as the Formlabs’ Clear Resin is UV-florescent, two 20W “Blacklight” UV fluorescent tubes were placed above the tank.

The sinking models were recorded using two Logitech C920 HD webcams (Logitech, Lausanne, Switzerland), placed at 90° to each other (SI Fig. 7) and recording at 960 pixels x 720 pixels and ∼30 frames per second, allowing monitoring of the position and orientation of the particle in 3D as it fell. As these consumer-grade webcams use a variable frame-rate system, a custom MATLAB script was used to initiate camera recording, recording both frames and frame timestamps. Videos were recorded for 500 frames (∼17 s). Sinking velocity was calculated over a central 0.8 m depth range, ensuring the model was at terminal velocity whilst also avoiding end effects which could slow the model as it reached the bottom of the tank. The curved walls of the tank introduced distortion, which was removed using the MATLAB toolbox “Camera Calibrator” (Mcandrew, 2004). Pixel size was 1.06 pixels per millimetre with a mean reprojection error of 0.5 pixels, therefore distance measurements (for calculating sinking velocities) were accurate to within 0.5 mm (0.06% of the traversed depth).

Models were tracked in distortion-corrected frames using a modified version of Trackbac (Guadayol et al., 2017; Guadayol, 2016). The per-frame centroid coordinates obtained were then paired with the timestamp values recorded to calculate average settling velocity components in 2D for each camera (below, *U*_*x*_ is horizontal speed from camera one, *U*_*y*_ is horizontal speed from camera two, and *U*_*z*,1_ and *U*_*z*,2_ are the vertical speeds corresponding to the two cameras). A resultant velocity magnitude was then calculated for each camera, and these two values averaged to yield a single estimate for *U* per experiment (SI Eqn 1).

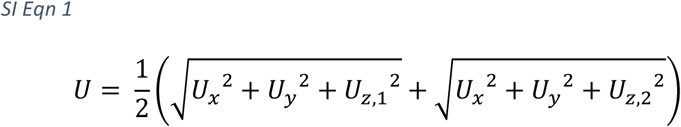

Each model was sunk five times and a mean *U* was calculated from these replicates. Replicates beyond a threshold of ±5% of the median sinking velocity were discarded from this average. Each model was dropped one additional time and photographed using a Canon 1200D DSLR camera (Tokyo, Japan) mounted on a tripod close to the tank, to obtain high resolution (18 megapixels) images which were used to determine model orientation (and thus *L* and *A*) during settling (SI Fig. 4).

**SI Fig. 6.**
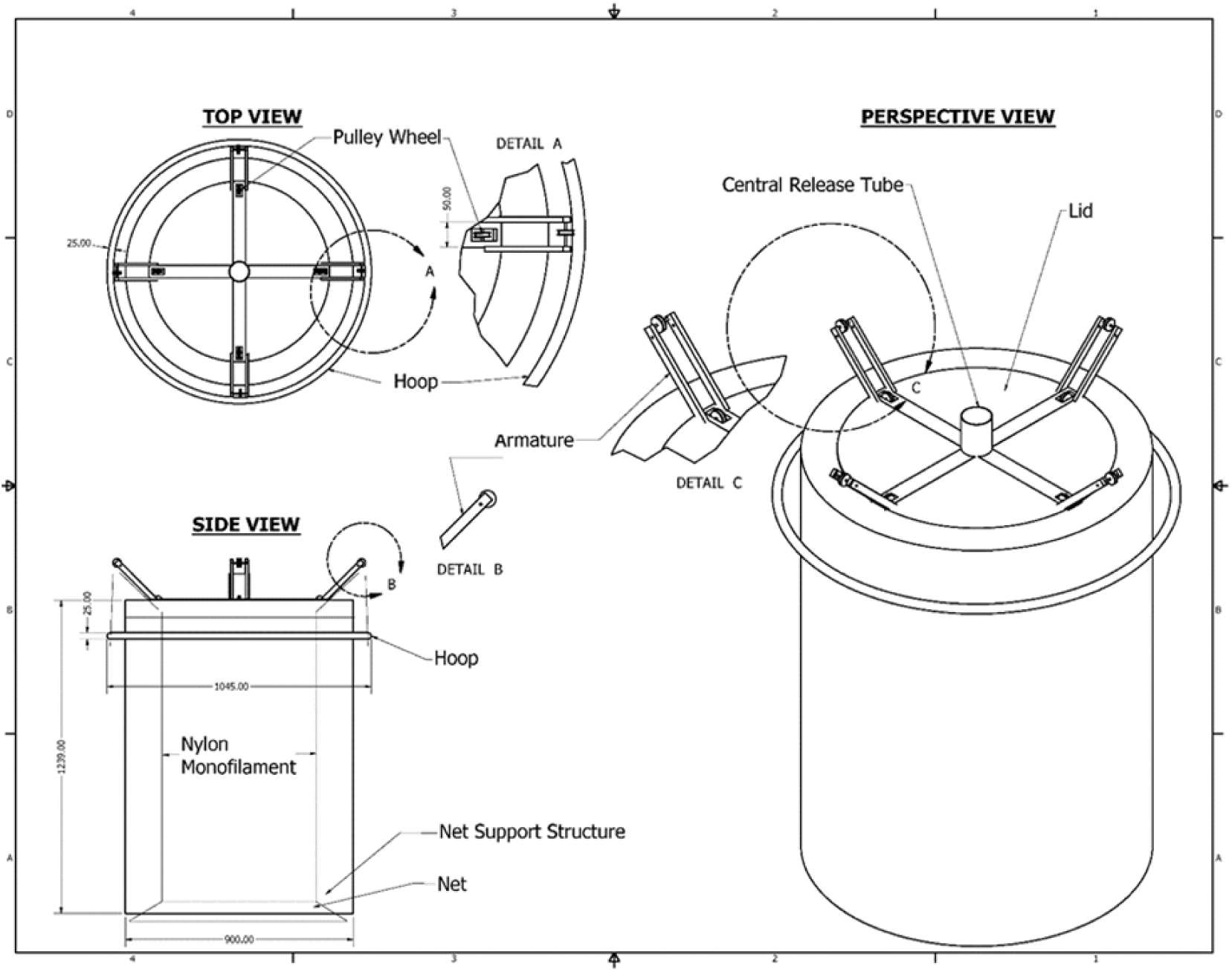
Schematic of the tank and model retrieval system. All measurements in mm.

**SI Fig. 7.**
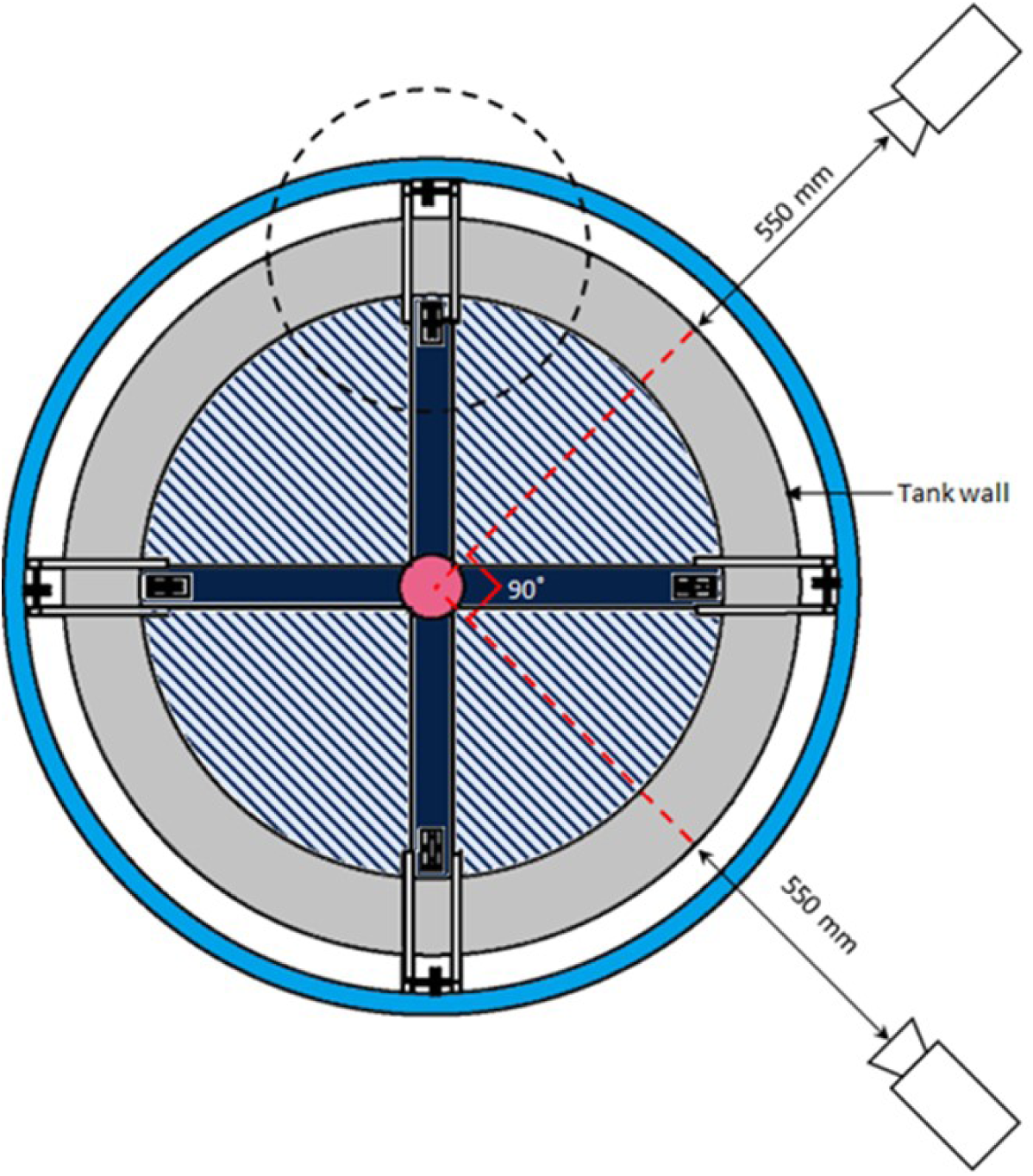
A top down view of the tank and camera positions. An external hoop (light blue) was used to raise the net from the bottom of the tank, via the system of pulleys and nylon monofilament mounted on the armatures (circled with black dashed line). The support arms (dark blue) held a central tube (pink) where the release mechanism was mounted. A ledge (grey) around the top of the tank was used to support the armatures, support arms and cover for when the tank was not in use. The open part of the tank is shown with dark blue hatching.

### SI 4. Further Applications

**SI Table 1.** A selection of fluids that could be used for enlarged, dynamically scaled models of settling particles. Air and water are provided as reference points; however, water could be used to study particles that normally operate in air. It should be noted that the density of the fluid needs to be lower than the density of the model in order for the model to sink. Fluids with custom viscosity can be created in principle by mixing various miscible fluids, e.g. water and glycerol or mineral and silicone oils.

| Fluid | $\rho_{fluid}$<br>(Kg m<sup>-3</sup>) | $\mu$<br>(Pa S) | Reference |
| --- | --- | --- | --- |
| Air | 1.225 | $1.85 \times 10^{-5}$ | (Munson et al., 1994) |
| Water | 1000 | 0.001 | (Munson et al., 1994) |
| Low Viscosity silicone oils | 918 | 0.004 | http://www.clearcoproducts.com/ |
| Mineral oils | 830 | 0.022 | This study |
| Medium Viscosity silicone oils | 960 | 0.048 | http://www.clearcoproducts.com/ |
| Glycerol/ Glycerine | 1260 | 0.943 | (Segur & Oderstar, 1951) |
| High Viscosity silicone oils | 975 | 4.875 | http://www.clearcoproducts.com/ |
| Super High Viscosity silicone oils | 979 | 293 | http://www.clearcoproducts.com/ |

**SI Table 2.**
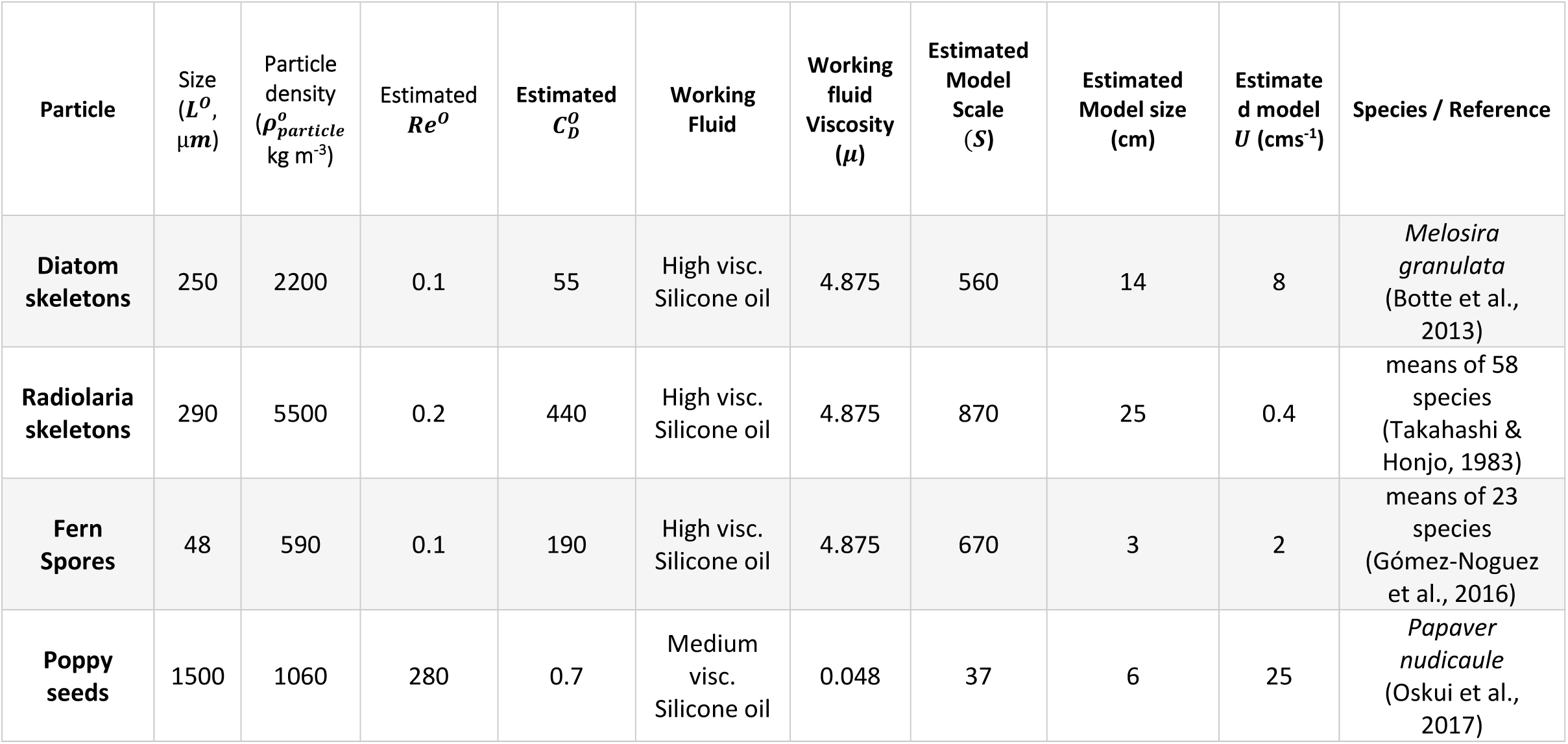
Examples of how one might implement our method to study settling of other biological particles. Typical particle sizes and densities are given, followed by estimates of Re^O^ and 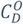 assuming filamentous diatom chains (with the theoretical C_D_(Re) relationship for cylinders at low Re, (Batchelor, 1959, Eq 4.10.16, p. 246)), and spherical particles for the other examples. If the proposed working fluids were used with 3D printed models similar to those used in our study (ρ_particle_ = 1120 kg m3), one would require models of the listed approximate scales and sizes, which would be expected to sink at the listed speeds.

## SI 5. Equation terms and Notations

*Re*: Reynolds number
*L*: Maximum length of particle parallel to the flow
*U*: Sinking speed of particle
*V*: particle volume
*A*: particle projected (frontal) area perpendicular to flow
*M*: particle mass
*ρ*_*fluid*_: fluid density
*ρ*_*particle*_: particle density
*μ*: fluid viscosity
*ΣF*: sum of external forces
*F*_*weight*_: particle weight
*F*_*drag*_: drag force
*F*_*buoyancy*_: buoyant force
*C*_*D*_: Coefficient of Drag
*Ψ*: 3D shape
*g*: acceleration due to gravity
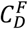: *C*_*D*_ determined through a force balance
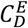: interpolating spline through (*Re, C*_*D*_) experimental data
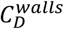: *C*_*D*_ with walls present (i.e. measured in the tank)
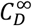: *C*_*D*_ in an unbounded domain (i.e. in the ocean)
*Ar*: Archimedes number
*S*: model scale factor
*o*: Value for real particle at natural operating point
*λ*: tank to particle diameter ratio
*K*: wall effects correction factor as per Happel and Brenner (1983)
*N*: number of data points
*H*: cubic spline interpolant for measured *V* vs *S*

## Acknowledgements

Our thanks go to Gregory Sutton and David Smith for their comments on a draft of this manuscript. We also thank Tatjana Hoehfurtner for segmentation of µCT scans and for her help with the 3D printing and cleaning of models. Additionally, the beam line scientists at PETRA III synchrotron storage ring at DESY in Germany for performing the high-resolution scans of foraminifera.

